# Peripheral kappa opioid receptor activation drives noxious cold hypersensitivity in mice

**DOI:** 10.1101/2020.10.04.325118

**Authors:** Manish K. Madasu, Loc V. Thang, Priyanka Chilukuri, Sree Palanisamy, Joel S. Arackal, Tayler D. Sheahan, Audra M. Foshage, Richard A. Houghten, Jay P. McLaughlin, Jordan G. McCall, Ream Al-Hasani

## Abstract

Noxious cold sensation is commonly associated with peripheral neuropathies, however, there has been limited progress in understanding the mechanism of cold pain. Here we identify a role for kappa opioid receptors (KOR) in driving noxious cold hypersensitivity. First, we show that systemic activation of KOR by the agonist U50,488 (U50), increases the latency to jump and the number of jumps on a cold plate at 3°C, and that the KOR antagonist NorBNI attenuates U50-induced noxious cold hypersensitivity. However, the central administration of NorBNI does not block U50-induced noxious cold hypersensitivity, suggesting that peripheral KOR may modulate this effect. To directly test this, we use the peripherally-restricted KOR agonist, ff(nle)r-NH2 and also show selective activation of peripheral KOR causes noxious cold hypersensitivity. To begin to understand how peripheral KOR drive noxious cold hypersensitivity we investigated whether KOR interact with transient receptor potential ankyrin 1(TRPA1) channels, known to facilitate the perception of noxious cold, in dorsal root ganglion (DRG). Using fluorescent *in situ* hybridization, we show that KOR mRNA colocalizes with the transcripts for the cold-activated TRPA1 channels in DRG. We also show a potentiation in intracellular calcium release in DRG neurons during the simultaneous application of the TRPA1 agonist, mustard oil (MO), and a KOR agonist, U50, when compared to MO alone. Together our data suggest that peripheral KOR may induce noxious cold hypersensitivity through modulation of TRPA1 channels.

## Introduction

Cold sensitivity is an elusive condition that has previously been defined as an exaggerated or abnormal reaction to cold exposure, causing discomfort or the avoidance of cold (Kay, 1985). Cold hypersensitivity is often associated with neuropathic pain from disorders such as multiple sclerosis, fibromyalgia, complex regional pain syndrome, and chemotherapy-induced peripheral neuropathy (Attal et al., 2009; Christogianni et al., 2018; Jensen and Finnerup, 2014; Tajerian and Clark, 2016; Wilbarger and Cook, 2011). 15% to 50% of neuropathic pain patients often experience heightened sensory abnormalities (Jensen & Finnerup, 2014). Medications used to treat neuropathic pain are predominantly non-steroidal anti-inflammatory drugs (NSAIDs), mu opioid agonists, and anti-epileptics (Kudel et al., 2019), all which have very limited success in relieving cold hypersensitivity.

While less in known about cold pain, significantly more is known about cold sensation, particularly how it is regulated by transient receptor potential (TRP) channels, TRPA1 and TRPM8 in DRG (Bautista et al., 2006a; Dhaka et al., 2008; McKemy et al., 2002a; Patapoutian et al., 2009; Pogorzala et al., 2013; Story et al., 2003). Most recently, it was shown that blocking K_v_1 potassium channels on large-diameter neurons in the periphery potentiated cold hypersensitivity in chronic pain states (MacDonald et al., 2021). Similarly loss of Nav1.9 sodium channel (Na_v_1.9^−/−^mice) known to predominantly express in DRG significantly attenuated chemotherapy-induced cold allodynia (Lolignier et al., 2015). These recent studies not only highlight the complex nature of cold-sensing mechanisms but also that this likely differs depending on the type of cold pain.

There is also some evidence implicating G protein-coupled receptors (GPCR) in sensing noxious, irritant, and inflammatory stimulants (Veldhuis et al., 2015) in tandem with TRP channels (Clapham, 2003). Specifically, activation of the kappa opioid receptor GPCR system is antinociceptive for noxious heat stimuli, analgesic in chronic pain models, inhibits itch transmission and drives pain-induced negative affect (Horan and Porreca, 1993; Liu et al., 2019; Massaly et al., 2019; Munanairi et al., 2018; Nguyen et al., 2021). Recent evidence suggests that centrally- and peripherally-expressed KOR modulates different behaviors. For example, our group and others have shown that central KOR activation and upregulation modulates negative affect associated with peripheral nerve injury and inflammatory pain models (Liu et al., 2019; Massaly et al., 2019). Conversely, recent studies suggest peripherally-restricted KOR agonists selectively inhibit chemical pain and mechanical hypersensitivity associated with capsaicin-induced neurogenic inflammatory pain model and a surgical incision model, respectively (Snyder et al., 2018). Together these findings highlight the complex role of the KOR system in different pain and sensation modalities.

Here we add another dimension to the complex role of KOR by uncovering a role for KOR in cold hypersensitivity in mice. We use pharmacological approaches to determine that peripheral KOR regulate cold hypersensitivity. We use *in situ* hybridization to show that KOR colocalize with TRPA1 and TRPM8 in the DRG. Finally, we use calcium imaging to demonstrate that KOR activation potentiates TRPA1-dependent calcium signaling in DRG neurons. In sum, we show that peripheral KOR activation enhances cold sensitivity and this behavior is likely mediated through potentiated TRPA1 activity.

## Methods

### Animals

Adult C57BL/6J male and female mice (25-30 g) were used for all the behavioral and *in situ* hybridization experiments. For calcium imaging experiments, we used TRPA1^−/−^ male and C57BL/6J male mice. All animals were 9 to 12 weeks at the beginning of the experiments. Mice were group-housed together with a 12/12 h dark/light cycle, given access to food pellets and water ad libitum (lights were turned on at 6:00 AM). Following weaning, all animals were transferred to a holding facility adjacent to the lab and acclimated to this animal facility for at least seven days before the experiments to minimize stress. Furthermore, all animals were acclimated to the behavior test room for at least two hours prior to each experiment. All procedures were approved by the Washington University in St. Louis Institutional Animal Care and Use Committee (IACUC) in accordance with the National Institutes of Health Guidelines for the Care and Use of Laboratory Animals.

### Drugs

U50,488 (U50, KOR agonist, 5 mg/kg i.p.) (CAS# 67197-96-0) and nor-Binaltorphimine dihydrochloride (norBNI, KOR antagonist, 10 mg/kg, i.p.) (CAS# 113158-34-2) were obtained from Tocris and dissolved in 0.9% sterile saline. Morphine sulfate (mu-opioid receptor (MOR) agonist, 10 mg/kg s.c.) procured from (Mckesson #996806) was dissolved in 0.9% sterile saline. CR845 analogues (ff(nle)r-NH2) (peripherally-restricted KOR agonist, 10 mg/kg, i.p.) were generated in the Mclaughlin lab (Alleyne et al, in revision) and dissolved 0.9% sterile saline.

### Hot/Cold Plate Assay

The hot/cold plate apparatus was adapted from the operant thermal plantar assay (Reker et al., 2020). The floor of the apparatus is made of one 12” × 12” × 1/4” aluminum plate (3003, MetalsDepot), fixed to a cold plate Peltier (Cold plate cooler, CP-061, TE Technology). The Peltier device is independently controlled by a power supply (PS-12–8, 4A, TE Technology) and temperature controller (TC-48–20, TE Technology). Long Cast Acrylic Tubing (15′ height, E-plastics) was used to contain the mice on the plate. Male and female C57BL/6J wildtype (WT) mice were habituated in plexiglass boxes for 30 mins prior to treatment with: 1) U50, KOR agonist, (5 mg/kg, i.p.) 2) norBNI, KOR antagonist (10 mg/kg, i.p.) 3) Saline (0.25ml) 4) CR845 peripherally-restricted KOR agonist (10 mg/kg i.p.). Following treatment, the mice were placed on the plate for 5 minutes, and the latency to jump and the number of jumps was recorded as nocifensive responses. The temperature of the plate varied from 3°C-42°C depending on the experiment. The plate’s temperature was continuously monitored with a visual temperature strip (Liquid crystal temperature indicating sheets-Telatemp) and a surface probe thermometer (Pro-surface thermapen-Thermoworks).

### Rectal temperature measurements

Core body temperature readings were obtained using a homeothermic monitoring system attached to a flexible rectal probe (Physitemp instrument Inc. TCAT2LV controller). On the test day, mice were left to acclimate in the test room for 90 minutes. Post-habituation mice were injected with saline (0.25ml) or U50 (5 mg/kg i.p.) or ff(nle)r-NH2 (10 mg/kg i.p.). Immediately post-injection, the mouse was hand-restrained and the tail lifted, and a lubricated probe gently inserted into the rectum to a fixed depth (typically, up to 1 cm) to obtain the rectal temperature. 30 minutes post-injection, the rectal temperatures were recorded. We made sure the depth of probe insertion is constant for each measurement.

### Paw surface temperature measurements

Paw surface temperature readings were obtained using FLIR One Pro LT (Android version). On the test day, mice were left to acclimate in the test room for 90 minutes. Post-habituation mice were injected with saline (0.25ml) or U50 (5 mg/kg i.p.). Immediately post-injection, the mouse was hand-restrained and then a thermal image was taken exposing the paws to obtain a clear reading of paw temperatures. 30 minutes post-injection, the paw surface temperatures were recorded. The images collected were processed on the FLIR one app (Windows10 64bit version) by identifying areas (sp1-5) on the paws to tabulate the paw surface temperatures.

### Tail withdrawal assay

Mice were habituated to the experimenter for a week prior to the assay and handled daily to minimize any stress that might impact the assay (Deuis et al., 2017). On experimental day, following U50 (5 mg/kg i.p.) or saline (0.25ml) injection, the mice were scruffed, and one-third of the distal end of the tail was immersed in the hot water bath set at 54.5°C. The time taken for the tail to twitch or flick was recorded. A single reading was recorded at 15, 30, 45, 60, 90, and 120 minutes following treatment.

### Von Frey test

Mechanical sensitivity was determined using manual von Frey filaments following U50 (5 mg/kg i.p.) or Saline (0.25ml) treatments. On the test day, mice were individually placed in an acrylic cylinder on a mesh floor and covered with a rectangular acrylic lid to prevent them from escaping. The mice were habituated to apparatus for 2 hours prior to testing. During the assay, a monofilament was applied perpendicularly to the plantar surface of the hind paw for 2–5 seconds. If the animal exhibited any nocifensive behaviors, including brisk paw withdrawal, licking, or shaking of the paw, either during or immediately following the stimulus, a response is recorded. In that case, such a response is considered positive (Deuis et al., 2017). We used the “ascending stimulus” method, applying monofilaments with increasing force until it elicits withdrawal response. The von Frey filament force that elicits this positive response represents the mechanical withdrawal threshold. The stimulus was repeated for both the paws (three readings from each paw) and their mean calculated. A 10-minute break was included between each reading to prevent hypersensitivity.

### Acetone Evaporation Test

The acetone evaporation test was used to measure aversive behaviors triggered by evaporative cooling (Deuis et al., 2017). On the test day, mice were left on the mesh floor for 90 minutes for acclimatization. Post-habituation, mice were injected with U50 (5 mg/kg i.p.) or saline (0.25ml) treatments. Post-injections, acetone (5–6 μl) was applied to the plantar surface of the hind paw using the open end of a blunt 1-ml syringe, eliciting a rapid paw withdrawal response (i.e., elevation, shaking, or licking). The acetone application was repeated for both the paws (three readings from each paw) and their mean was calculated. The response was recorded for one minute following each acetone application. A 10-minute break was included between each reading to prevent hypersensitivity (Colburn et al., 2007; Slivicki et al., 2018). Sensitivity to cold is recorded either by quantifying paw directed behavior, or scoring the severity of the response (for example: 0, no response; 1, brisk withdrawal or flick of the paw; 2, repeated flicking of the paw; 3, repeated flicking of the hind paw and licking of the paw) or latency to lick the paw.

### Open field test (OFT)

The OFT apparatus is a 2,500 cm^2^ arena, in which the mice were free to explore for 20 minutes. 50% of the total OFT area was defined as the center (Al-Hasani et al., 2015; McCall et al., 2015). Lighting was stabilized at ~25 lux for anxiety-like behaviors. To determine sedative and anxiety-like behavior, distance moved in the apparatus and the time spent exploring the center were quantified, respectively. On the test day, mice were left in the OFT room for 90 mins for acclimatization to the environment. Post-habituation mice were injected with saline or U50, placed in the OFT arena, and behavior was recorded for 20 minutes. Movements were video recorded and analyzed using Ethovision 13 (Noldus Information Technologies).

### Intracerebroventricular injections

Mice received intracerebroventricular (ICV) injections as previously described (Bruchas et al., 2009). Briefly, mice were anesthetized in an induction chamber (3% Isoflurane) and placed into a stereotaxic frame (Kopf Instruments, Model 942), where they were maintained at 2–2.5% isoflurane. A craniotomy was performed unilaterally into the lateral ventricle with either NorBNI (30μmol, 2μl) or artificial cerebrospinal fluid (aCSF) at 200 nl/min for 10 mins using a beveled Hamilton syringe (10 μL-701 N with beveled tip) (stereotaxic coordinates from Bregma: A/P: +0.40mm, M/L: +1.5mm, D/V: −3mm), to block central KOR (Bruchas et al., 2009; Shirayama et al., 2004). The skin was sutured after the injection using sterile nylon sutures (6.0 mm), and mice were allowed to recover for a week before any behavioral experiments. On the experimental day, post-surgery mice were systemically injected with either U50 (5 mg/kg i.p.) or saline (0.25ml) and were exposed to the cold plate. The ICV injection placements were confirmed using immunohistochemistry.

### Immunohistochemistry

Histological verification of the injection site was performed as described (Al-Hasani et al., 2013). Mice were briefly anesthetized with 0.2ml cocktail (ketamine (100 mg/ml), xylazine (20 mg/ml) and acepromazine (10 mg/ml) and intracardially perfused with ice-cold 4% paraformaldehyde in phosphate buffer (PB). Brains were dissected, post-fixed 24 h at 4 °C and cryoprotected with 30% sucrose solution in 0.1 M PB at 4 °C for at least 24 h, cut into 30-μm sections, and processed for immunostaining. The ICV injection placements were confirmed using a Leica fluorescent microscope (DM6 series scope) with the Paxinos-Watson atlas (Paxinos, G. and Watson, 1998) as a reference. In total, n=76 out of 85 WT mice (male and female) had the injections landing in the lateral ventricle and the rest were excluded from the analysis.

### *In situ* Hybridization

Following rapid decapitation of C57BL6/J mice, DRG were rapidly frozen on dry ice in the mounting media, and then the tissue harvested was stored at −80°C. DRG sections were cut at 5-7 μM at −20°C and thaw-mounted onto Super Frost Plus slides (Fisher, Waltham, MA). Slides were stored at −80°C until the following day. Fluorescent *in situ* hybridization (ISH) was performed according to the RNAScope 2.0 Fluorescent Multiple Kit User Manual for Fresh Frozen Tissue (Advanced Cell Diagnostics, Inc.), as described (Wang et al., 2012). Briefly, sections were fixed in 4% PFA, dehydrated with alcohol (50%, 75%, 100%) concentrations in the respective order in accordance with the protocol. Sections were pretreated with hydrogen peroxide for 15 min at room temperature. Following this, the sections were washed in the 1X PBS solution twice for 2 min each. Post-wash, the sections were pretreated with protease IV solution. Sections were then incubated for target probes for mouse *trpa1*(400211) (C1), *trpm8 (*420451) (C2), *oprk1* (316111) (C3). Probes were obtained from Advanced Cell Diagnostics. Following probe hybridization, sections underwent a series of probe signal amplification steps followed by incubation with fluorescently labeled probes designed to target the specified channel associated with *trpa1* (fluorescein), *trpm8* (cyanin3), *oprk1* (cyanin 5). Slides were counterstained with DAPI, and coverslips were mounted with Vectashield Hard Set mounting medium (Vector Laboratories). Images were obtained on a Leica fluorescent microscope, and the expression was quantified manually using the Leica DM6 series scope by a blinded experimenter. DRGs were imaged on a Leica fluorescent microscope (DM6 series scope) at 5X, 20X, and 40X magnification. 2-3 images were acquired of each mouse DRG section, and 4-5 DRGs were imaged per mouse (n=4 for male, n=2 for females). Total cell counts for the section were assessed by counting all of the DAPI in the DRG section. In the ISH assay, each punctate dot represents a single target mRNA molecule. To avoid false positives, we set a threshold of a minimum of two puncta expressing cells only. The target genes expression was quantified manually by counting the DAPI cells expressing the puncta. The quantified expression was averaged within a sample, across mice and expressed as a pie chart. Each *oprk1* positive cell was assessed for colocalization with *trpm8* and *trpa1* using 40X magnification.

### Mouse DRG cultures

10-week-old male and female mice were euthanized under isoflurane by decapitation and lumbar DRG were removed (Sheahan et al., 2018; Sleigh et al., 2016). DRG culture media was prepared fresh using Neurobasal A medium (Invitrogen) with 100 U/mL penicillin/streptomycin (Corning), 2 mm GlutaMAX (Life Technologies), 2% B27 (Gibco), and 5% fetal bovine serum (Gibco). DRG were incubated in papain (40 U, Worthington) for 20 min at 37°C, supplemented with 5% CO2. DRG were then rinsed and incubated in collagenase (Sigma-Aldrich) for 20 min, following which they were manually triturated with Pasteur pipettes to dissociate neurons, passed through a sieve of 40-μm filter, and plated onto collagen (Sigma-Aldrich)-coated 12-mm glass coverslips (Thermo Fisher Scientific). Neurons were maintained in culture media for two days prior to calcium imaging experiments.

### Calcium Imaging

To determine calcium dynamics, cultured DRG neurons from C57BL/6J male and female, TRPA1^−/−^ male mice were loaded with Fura-2 (3 μg/mL, Life Technologies) and pluronic acid (1:1) for 45-60 min to enable visualization of changes in calcium concentrations (Munanairi et al., 2018; Snyder et al., 2018). Neurons were then incubated in Tyrode’s solution for 15-20 min to allow for de-esterification of Fura-2 AM. Tyrode’s solution consisted of (in mm): 130 NaCl, 5 KCl, 2 CaCl2, 1 MgCl2, 30 glucose, and 10 Hepes and was made fresh on the experimental day. On the day of recording, coverslips were placed into a temperature-controlled chamber (37°C) with Tyrode’s solution. The cultures were treated with either KOR agonist (U50488, 10 μM) alone, TRPA1 agonist (Mustard oil (MO) 100μM), or a combination of both. The change in the intensity of the Ca^2+^indicator is quantified to estimate the change in the concentration of free Ca^2+^ using calcium imaging software suite (Leica Systems) to record fluorescence emission at alternating excitation wavelengths of 357 and 380 nm. A KCl response at the end of an experiment served as a positive control for cell health and activity. Detailed timelines are included in the figure sets.

### Statistical Analysis

All data samples were tested for homogeneity of variance and normality before being assigned to any parametric analysis. All experiments were performed in multiple cohorts, including all treatment groups in each round, to avoid any unspecific day/condition effect. Treatments were randomly assigned to animals before testing. G*Power was used to estimate effect sizes and to compute power analyses. Statistical significance was considered \**p*< 0.05, \*\**p* < 0.01, \*\*\**p*< 0.001, and \*\*\*\**p*< 0.0001, as determined by parametric or nonparametric analysis. Parametric data were analyzed using two-way ANOVA repeated measures for tail withdrawal assay analysis, two-way ANOVA for mechanical sensitivity analysis. For ordinal ranking such as scoring the number of jumps (nonparametric data), Kruskal-Wallis followed by the Dunn’s multiple comparison test for analyzing the cold plate assay data. For the temperature curve data, nonparametric (two-way ANOVA) mixed-effects analysis was followed by Sidak’s *post-hoc* test for the cold plate assay analysis. Mann Whitney U test for cold plate data using peripheral novel agonist. Statistical analyses were performed in GraphPad Prism 8.0. All data are expressed as mean ± SEM.

## Results

### Activation of KOR induces noxious cold hypersensitivity

To determine whether activation of KOR mediates temperature-dependent hypersensitivity, we recorded jumping behavior across a range of temperatures (3°C, 10°C, 15°C, 20°C, 30°C, 42°C) (**Fig 1A**). Jumping behavior in this assay has been well-established as a measure of nocifensive behavior (Allchorne et al., 2005; Castellanos et al., 2020; Deuis et al., 2017). We showed that the KOR agonist, U50 (5 mg/kg i.p.) significantly increased the number of jumps when compared to controls at 3°C in males and females (**Fig 1B&C**). This jumping behavior was supported by a significant decrease in latency to jump in male and females at 3 °C (**Fig 1D&E**). Together, these results show that KOR activation selectively induces hypersensitivity to noxious cold. An increase in the number of jumps was also observed at 42°C, but this effect also occurs in the saline-treated groups, and the U50-treated male mice suggesting that this was not due to activation of KOR, but rather the noxious hot temperature itself (Yalcin et al., 2009) **(Fig 1C**).

**Figure 1:**
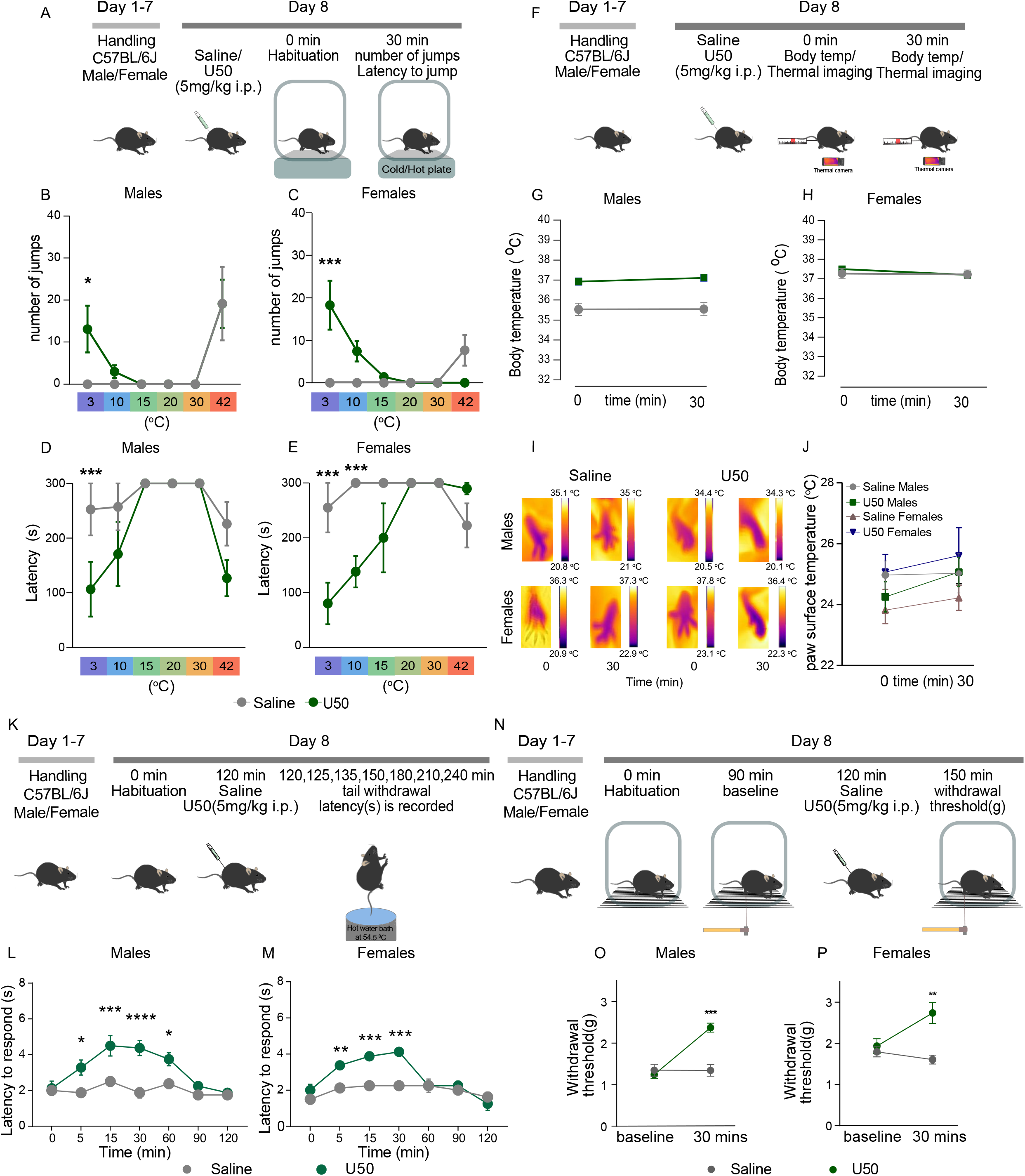
KOR-induces hypersensitivity at 3°C A) Outline of the experimental procedure. WT mice were injected with either saline or U50 (KOR agonist, 5 mg/kg, i.p.) 7 days post-handling. Post injection mice were exposed to cold/hot plate assay across a range of temperatures 3°C, 10°C, 15°C, 20°C, 30°C, 42°C. B) In males, KOR agonist U50 (5mg/kg i.p.) significantly increased the number of jumps compared to controls at 3°C. Nonparametric mixed effects analysis (two-way ANOVA) revealed temperature effect *F* _(5, 52)_ = 8.92 followed by Sidak’s *post-hoc* test (\**P*<0.05 vs. respective saline treatment). Data are presented as mean ± SEM; n = 5-7/group. C) In females, KOR agonist U50 (5mg/kg i.p.) significantly increased the number of jumps compared to controls at 3°C. Nonparametric mixed effects analysis (two-way ANOVA) revealed temperature effect *F* _(5, 57)_ = 3.937, treatment effect *F* _(1, 57)_ = 4.956 and a treatment x temperature *F* _(5, 57)_ = 6.191 followed by Sidak’s *post-hoc* test (\*\*\**P*<0.001 vs respective saline treatment). Data are presented as mean ± SEM; n = 5-7/group. D) Latency to jump following KOR activation is decreased in males at 3°C only. Nonparametric mixed effects analysis (two-way ANOVA) revealed temperature effect *F* _(5, 53)_ = 4.792, treatment effect *F* _(1, 53)_ = 8.665 and a treatment x temperature *F* _(5, 53)_ = 2.446 followed by Sidak’s *post-hoc* test (\*\*\**P*<0.001 vs respective saline treatment). Data are presented as mean ± SEM; n = 5-7/group. E) Latency to jump following KOR activation is decreased in females at 3°C and 10°C; Nonparametric mixed effects analysis (two-way ANOVA) revealed temperature effect *F* _(5, 30)_ = 5.917, treatment effect *F* _(1, 29)_ = 14.82 and a treatment x temperature *F* _(5, 29)_ = 6.722 followed by Sidak’s *post-hoc* test (\*\*\**P*<0.001 vs respective saline treatment). Data are presented as mean ± SEM; n = 5-7/group F) Outline of the experimental procedure. Timeline of the assay, wherein mice are habituated, baseline rectal temperature is measured along with thermal imaging of paw surface temperatures, followed by systemic injections of saline or U50 and post-treatment temperatures were recorded. G) KOR agonist U50 did not alter core body temperature 30 min post drug administration in males. Data are expressed as mean±SEM; n=6-10/group. H) KOR agonist U50 did not affect core body temperature 30 mins post drug administration in females. Data are expressed as mean±SEM; n=6-10/group. I) Representative infrared images from right hind paws of a male and female mouse using FLIR infrared camera pre-and post-saline/U50 adminstration. J) KOR agonist U50 did not alter paw surface temperatures in males and females. Data are expressed as mean±SEM; n=4/group. K) Outline of experimental procedure, WT mice were injected with either saline or U50 (KOR agonist, 5 mg/kg, i.p.) 7 days post-handling. Post injection mice were exposed to tail withdrawal assay at 0 mins, 15 mins, 30 mins, 60 mins, 90 mins and 120 mins. L) U50-mediated antinociception in males, parametric repeated measures ANOVA (two-way ANOVA) revealed time effect *F* _(6, 97)_ = 4.054, dose effect *F* _(6, 97)_ = 8.036 and time x dose effect *F* _(1, 97)_ = 42.85 followed by Sidak’s *post-hoc* test (\**P*<0.05, \*\**P*<0.01 \*\*\**P*<0.001 vs respective saline treatment). Data are expressed as mean± SEM; n=7-8/group. M) U50-mediated antinociception in females, parametric repeated measures ANOVA (two-way ANOVA) revealed time effect *F* _(6, 98)_ = 5.620, dose effect *F* _(6, 98)_ = 13.33 and time x dose effect *F* _(1, 98)_ = 28.63 followed by Sidak’s *post-hoc* test (\*\**P*<0.01 \*\*\**P*<0.001 vs respective saline treatment). Data are expressed as mean± SEM; n=7-8/group. N) Timeline of experimental procedures for von Frey assay, mice were habituated and baseline withdrawal threshold was scored, followed by systemic injection of saline or U50 (KOR agonist, 5 mg/kg, i.p.). 30 mins post-treatment response to mechanical stimulus was again recorded. O) U50 blocked mechanical sensitivity in males, parametric repeated measures ANOVA (two-way ANOVA) revealed time effect *F* _(1, 15)_ = 36, dose-effect *F* _(1, 15)_ = 10.96 and time x dose-effect *F* _(1, 15)_ = 36.54 followed by Sidak’s *post-hoc* test (\*\*\**P*<0.001 vs. respective saline treatment). Data are expressed as mean± SEM; n=6-8/group. P) U50 blocked mechanical sensitivity in females, parametric repeated measures ANOVA (two-way ANOVA) revealed dose-effect *F* _(1, 12)_ = 15.26 and time X dose-effect *F* _(1, 12)_ = 5.127 followed by Sidak’s *post-hoc* test \*\**P*<0.001 vs. respective saline treatment at 30 mins). Data are expressed as mean± SEM; n=6-8/group.

We demonstrated that U50 does not alter core body temperature (**Fig 1F,G&H**) or paw surface temperature (**Fig 1&J**) in males and females, as compared to saline controls, using rectal probe and paw surface temperature using thermal imaging. This suggested that KOR-mediated cold hypersensitivity effects were not due to central thermoregulation. We also corroborated published findings showing that U50 (5 mg/kg i.p.) was antinociceptive in both the warm water tail-withdrawal assay (**Fig 1K,L&M**) and the von Frey test of mechanical sensitivity (**Fig 1O&P**). To further substantiate that KOR-induced noxious cold hypersensitivity was temperature-dependent, we used the acetone evaporation test. Acetone is known to mimic cool temperatures in the range of 15–21°C, but not noxious cold temperatures (Colburn et al., 2007; Leith et al., 2010). In this test, we showed that U50-mediated KOR activation did not alter paw withdrawal behavior (**Fig S1B**), or duration of nocifensive responses (**Fig S1C**). Importantly, we showed that the dose of U50 used in this study (5 mg/kg i.p) did not have sedative effect (common at higher doses), as demonstrated by no differences in locomotor activity between the groups (**Fig S1E–G**).

### Potentiation of noxious cold hypersensitivity is KOR selective

To determine the necessity of KOR’s role in cold hypersensitivity at 3°C, we pharmacologically blocked KOR using norBNI (KOR antagonist, 10 mg/kg, i.p.), which blocked U50-induced noxious cold hypersensitivity on the cold plate, in both male and female mice (**Fig 2A–C**). Furthermore, norBNI (i.p.) alone did not alter any nocifensive response irrespective of the sex (**Fig B&C**). We also showed that activation of mu opioid receptors with morphine (10 mg/kg s.c.), did not drive noxious cold hypersensitivity in male and female mice (**Fig 2E&F**), suggesting that this a KOR-selective effect. Together, these data suggest that KOR potentiates cold hypersensitivity at noxious cold temperatures (3°C).

**Figure 2:**
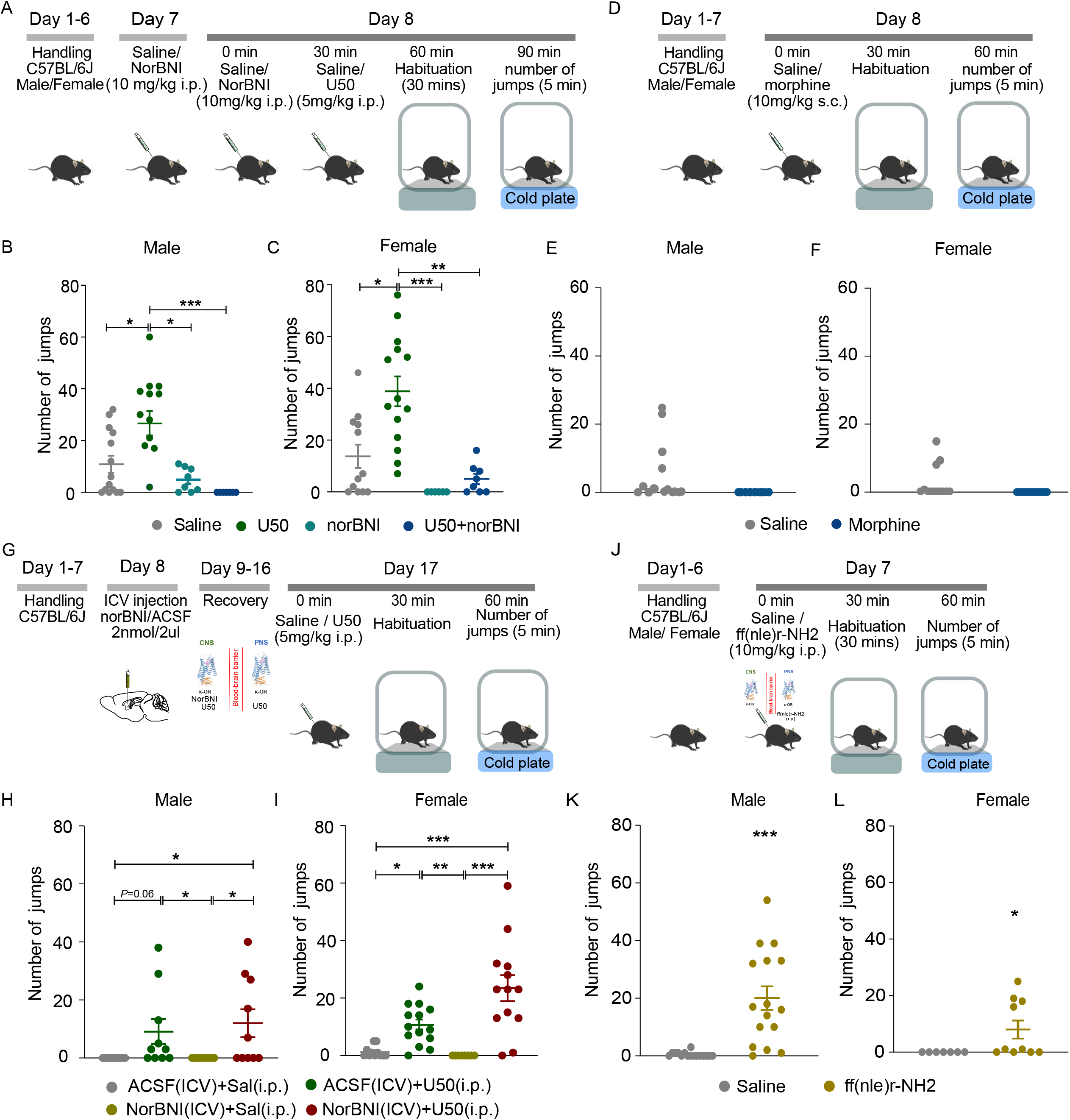
Potentiation of noxious cold hypersensitivity is KOR selective A) Outline of the experimental procedure. Timeline of the systemic injections of saline, U50, or norBNI, followed by habituation and cold plate assay at 3°C. B) In males, activation of KOR by U50 significantly increased the number of jumps on a 3°C cold plate. Kruskal-Wallis test revealed treatment effect *H* = 23.35 followed by Dunn’s *post-hoc* test (U50 vs. Saline \**P*<0.05). KOR antagonism with norBNI blocked cold hypersensitivity on a cold plate at 3°C (U50 vs U50+norBNI \*\*\**P*<0.001). Data are presented as mean±SEM n=8-14/group. C) In females, activation of KOR by U50 significantly increased the number of jumps on a 3°C cold plate. Kruskal-Wallis test revealed treatment effect *H* = 23.28 followed by Dunn’s *post-hoc* analysis (U50 vs. Saline \**P*<0.05). KOR antagonism with norBNI blocked cold hypersensitivity on a cold plate at 3°C (U50 vs U50+norBNI \*\**P*<0.01). Data are presented as mean±SEM n=8-14/group. D) Outline of the experimental procedure. Timeline of the systemic injections of morphine, followed by habituation, and cold plate assay at 3°C. E) Morphine did not affect cold sensitivity in males. Data are expressed as mean±SEM; n=10-12/group. F) Morphine did not affect cold sensitivity in females. Data are expressed as mean±SEM; n=10-12/group. G) Outline of the experimental procedure. Timeline of the ICV injections (NorBNI or aCSF), followed by habituation, systemic U50, or saline injections followed by cold plate assay at 3°C. H) In males, mice infused with aCSF (ICV) and injected with U50 (i.p.) showed cold hypersensitivity, as compared to the control group on 3°C cold plate assay. Kruskal-Wallis test revealed treatment effect *H* = 12.45 followed by Dunn’s *post-hoc* analysis. (aCSF (ICV)+U50 (i.p.) vs aCSF (ICV)+saline(i.p.) *P*=0.06). Central KOR antagonism with norBNI had no effect on cold hypersensitivity in males on a cold plate at 3°C (aCSF (ICV)+saline (i.p.) vs. norBNI (ICV)+U50 (i.p.) \**P*<0.05). Central KOR antagonism without peripheral activation has shown no cold hypersensitivity on a cold plate at 3°C (NorBNI (ICV)+saline (i.p.) vs. norBNI (ICV)+U50 (i.p.) \**P*<0.05). Data are presented as mean±SEM n=8-14/group. I) In females, mice infused with aCSF (ICV) and U50 (i.p.) showed cold hypersensitivity, as compared to the control group on 3°C cold plate assay. Kruskal-Wallis test revealed treatment effect *H* = 28.67 followed by Dunn’s *post-hoc* analysis (aCSF (ICV)+U50 (i.p.) vs aCSF (ICV)+saline(i.p.) \**P*<0.05). Central KOR antagonism with norBNI, had no effect on cold hypersensitivity in females on a cold plate at 3°C (aCSF (ICV)+saline (i.p.) vs. NorBNI (ICV)+U50(i.p.) \*\*\**P*<0.01). Central KOR antagonism without peripheral activation has shown no cold hypersensitivity on a cold plate at 3°C (norBNI (ICV)+saline (i.p.) vs. NorBNI (ICV)+U50(i.p.) \*\**P*<0.01). Data are presented as mean±SEM n=8-14/group. J) Outline of the experimental procedure. Timeline of the systemic injections of saline or ffir-NH2 (10mg/kg i.p.), followed by habituation and cold plate assay at 3°C. (K) Peripherally restricted agonist ffir-NH2 (10mg/kg i.p.) significantly increased jumping on the cold plate at 3°C, compared to controls, in males. Mann-Whitney U test revealed a treatment effect (Males-saline (i.p.) vs. ff(nlr)r-NH2 (i.p.) \*\*\**P*<0.001). Data expressed as mean±SEM n=7-16/group. L) Peripherally restricted agonist ffir-NH2 (10mg/kg i.p.) significantly increased jumping on the cold plate at 3°C, compared to controls, in females. Mann-Whitney U test revealed a treatment effect (females-saline (i.p.) vs. ff(nlr)r-NH2 (i.p.) \**P*<0.05). Data are presented as mean±SEM n=7-16/group.

### Peripheral KOR mediates noxious cold hypersensitivity

To determine whether KOR expressed centrally and/or peripherally mediates the observed increase in noxious cold hypersensitivity, we first blocked central KOR by injecting norBNI intracerebroventricularly (ICV) and administered U50 systemically (i.p.) before placing the mice on the cold plate (**Fig 2G**). Central administration of norBNI (norBNI, ICV + U50, i.p.) did not attenuate or block the observed U50-induced noxious cold hypersensitivity, as compared to controls (aCSF, ICV + U50, i.p.) in both males and females (**Fig 2H&I**). Central norBNI administration alone did not alter cold sensitivity, as compared to the controls (aCSF, ICV+ saline, i.p.) (**Fig 2H&I**). Mice infused with aCSF, ICV and U50, i.p. showed increased cold sensitivity, as compared to the control group (aCSF, ICV+ saline, i.p.), in both male and female mice (**Fig 2H&I**). Together these data show that central KOR does not seem to mediate noxious cold hypersensitivity. To extend these findings and directly address whether the KOR-induced noxious cold hypersensitivity is modulated by peripherally-expressed KOR, we used a peripherally restricted KOR agonist ff(nle)r-NH2 (10 mg/kg i.p.) (Alleyne et al, in revision). We showed that ff(nle)r-NH2 significantly increased jumping on the cold plate at 3°C, compared to controls, in both males and females (**Fig 2K&L**). We also demonstrated that ff(nle)r-NH2 did not alter core body temperature (**Fig S2A–C**) or paw surface temperature (**Fig S2D&E**), as compared to saline controls, using rectal probe and thermal imaging, respectively. Together, these data suggest that peripherally expressed KOR induces noxious cold hypersensitivity.

### KOR colocalizes with trpa1 and trpm8 transcripts and potentiates Ca^2+^ mobilization via TRPA1 in DRG

To further investigate how peripheral KOR mediates cold hypersensitivity we explored TRP channels, known to be necessary for temperature sensation. Specifically, the TRPA1 and TRPM8 channels have been most widely associated with cold sensitivity (MacDonald et al., 2021; McKemy et al., 2002b; Patapoutian et al., 2009). To determine whether KOR were expressed in TRPA1- and/or TRPM8-expressing cells in DRG, we performed *in situ* hybridization against the mRNA for KOR (*oprk1*), TRPA1 (*trpa1*), and TRPM8 (*trpm8*) in male (**Fig 3A–F**) and female (**Fig 3G–L**). We found that in male DRGs, 6.01% of all DRG cells expressed *oprk1* (**Fig 3M**), 17% expressed *trpa1* (**Fig 3N**), and 5.2% of the cells expressed *trpm8* (**Fig 3O**). 2% of the cells expressing *oprk1* colocalized with *trpa1* positive cells (**Fig 3P**), 2.7 % with *trpm8* (**Fig 3Q**) and 0.1 % with both the *trpa1* and *trpm8* (**Fig 3R**). The coexpression of *oprk1* with *trpm8* transcripts was higher in male DRGs when compared to female DRGs (**Fig 3Q**). In female DRGs, 6.8% of all DRG cells expressed *oprk1* (**Fig 3M**), 20% expressed *trpa1* (**Fig 3N**), and 5.25% expressed *trpm8* (**Fig 3O**). 2.8% of the cells expressing *oprk1* colocalized with *trpa1* positive cells (**Fig 3P**), 0.747% with *trpm8* (**Fig 3Q**), which was significantly different when compared to males and 0.3% cells expressed both the *trpa1* and *trpm8* (**Fig 3S**). When closely examining the proportion of cells expressing *oprk1* transcripts (**Fig 3T&U**), we show that a 45% of *oprk1* expressing cells in the DRG also express *trpa1* in male and female mice. Furthermore 13% of *oprk1* expressing cells in the DRG also express *trpm8* in male and female mice (**Fig 3T&U**). Together, both the TRP channels colocalization account to 60% of *oprk1* cell population in male and female mice (**Fig 3T&U**). The level of expression was similar between males and females except for *trpm8+oprk1* colocalization.

**Figure 3:**
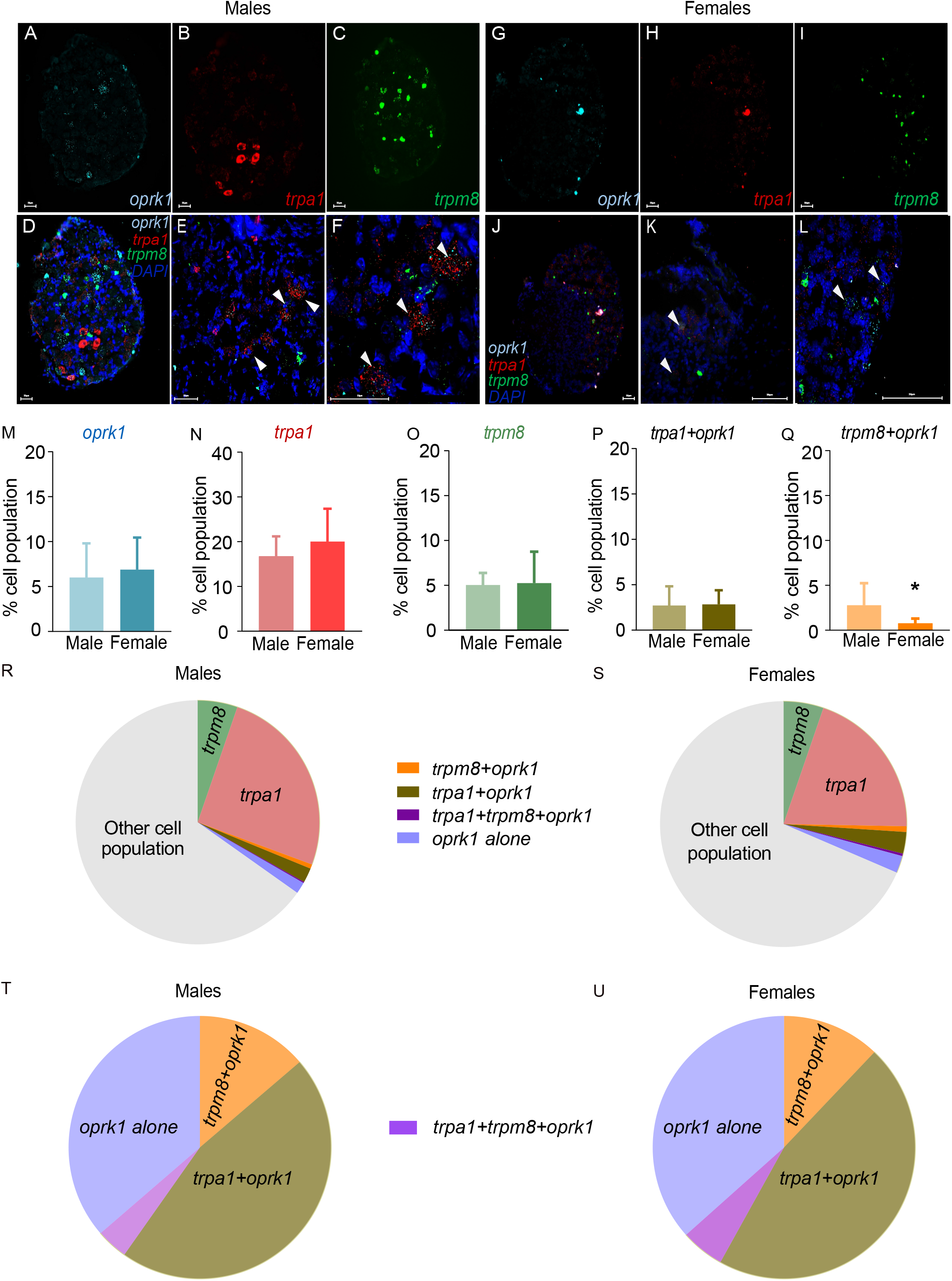
Transcripts for KOR, TRPA1, and TRPM8 are differentially co-localized in mouse DRG A) Representative image of Fluorescent *in situ* hybridization in a DRG section for *oprk1* in male mice, shown in blue. B) Representative image of Fluorescent *in situ* hybridization in a DRG section for *trpa1* in male mice, shown in red. C) Representative image of Fluorescent *in situ* hybridization in a DRG section for *trpm8 transcripts* in male mice, shown in green. D) Representative image of Fluorescent *in situ* hybridization in a DRG section of all the three transcripts *oprk1, trpa1, trpm8* and DAPI in male mice. E) Representative image of Fluorescent *in situ* hybridization in a DRG section showing the colocalization of *oprk1* with *trpm8* and *trpa1* at 40x in male mice. F) Representative image of Fluorescent *in situ* hybridization in a DRG section showing the colocalization of *oprk1* with *trpm8* and *trpa1* at 100x in male mice. G) Representative image of Fluorescent *in situ* hybridization in a DRG section for *oprk1* in female mice, shown in blue. H) Representative image of Fluorescent *in situ* hybridization in a DRG section for *trpa1* in female mice, shown in read. I) Representative image of Fluorescent *in situ* hybridization in a DRG section for *trpm8 transcripts* in female mice, shown in green. J) Representative image of Fluorescent *in situ* hybridization in a DRG section of all three transcripts and DAPI in female mice. K) Representative image of Fluorescent *in situ* hybridization in a DRG section showing colocalization of *oprk1* with *trpm8* and *trpa1* at 40x in female mice. L) Representative image of Fluorescent *in situ* hybridization in a DRG section showing colocalization of *oprk1* with *trpm8* and *trpa1* at 100x in female mice. M) Expression of *oprk1* transcripts in female mice were similar to the male mice in the DRG. Data are presented as mean±SEM; n=4/group. N) Expression of *trpa1* transcripts in female mice were similar to the male mice in the DRG. Data are presented as mean±SEM; n=4/group. O) Expression of *trpm8* transcripts in female mice were similar to the male mice in the DRG. Data are presented as mean±SEM; n=4/group. P) No difference in colocalization of *oprk1+trpa1* transcripts in DRG between males and females. Data are presented as mean±SEM; n=4/group. Q) Colocalization of *oprk1+trpm8* transcripts is higher in males compared to the females in the DRGs. Independent *t*-test analysis revealed a significant difference between males vs. females *t*_*6*_ = 0.801 \**P*< 0.05. Data are presented as mean±SEM; n=4/group. R) Quantification of *oprk1*, *trpm8, trpa1,* and co-expression of *trpa1* and *trpm8* transcripts with *oprk1* in male DRG. Data are presented as mean±SEM; n=4/group. S) Quantification of *oprk1*, *trpm8, trpa1,* and co-expression of *trpa1* and *trpm8* transcripts with *oprk1* in female DRG. Data are presented as mean±SEM; n=4/group. T) Quantification of co-expression of *trpa1, trpm8* or both transcripts with *oprk1* in male DRG. Data are presented as mean±SEM; n=4/group. U) Quantification of co-expression of *trpa1, trpm8* or both transcripts with *oprk1* in female DRG. Data are presented as mean±SEM; n=4/group.

To understand how peripheral KOR might induce noxious cold hypersensitivity, we measured calcium responses in cultured DRG following simultaneous application of MO (TRPA1 agonist) and U50 (**Fig 4A–C**) in male WT mice. U50 together with MO, significantly increased Ca^2+^ mobilization when compared to MO alone in DRG neurons in WT male mice (**Fig 4 B–D**). Application of U50 alone had no effect on mobilization of Ca^2+^ when compared to MO or MO+U50 treatments in WT male mice (**Fig 4D**). In TRPA1 knockout mice (TRPA1^−/−^), MO application alone and together with U50 had no effect on Ca^+2^ signaling (**Fig 4E–G**). Together our data suggests that KOR-induced noxious cold hypersensitivity is mediated through enhanced activity at TRPA1 channels.

**Figure 4:**
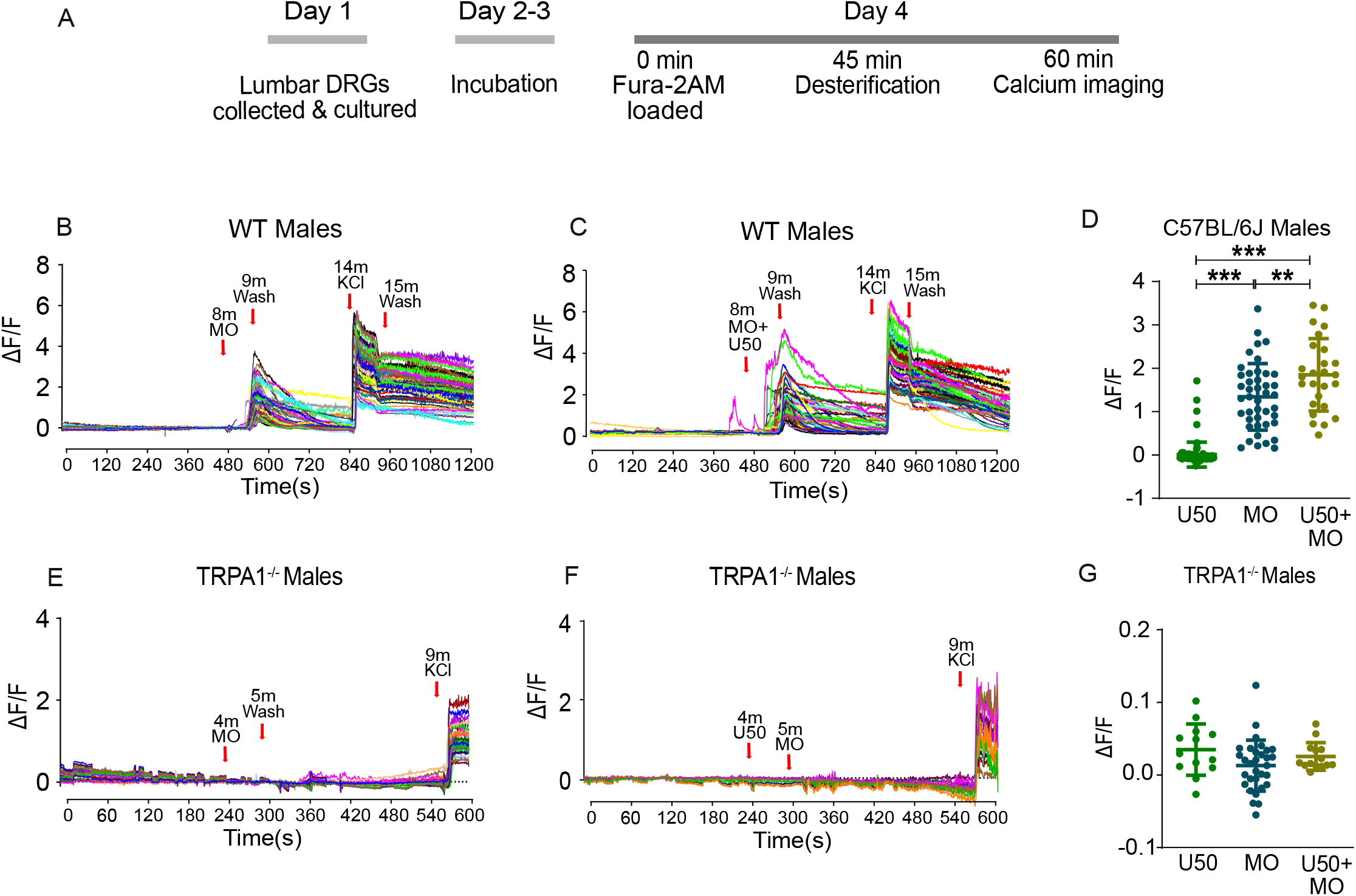
KOR activation potentiates TRPA1-dependent calcium signaling A) Outline of the experimental procedure. Timeline of the dissection of lumbar DRG, followed by culturing and calcium imaging of the DRG neurons. B) Representative traces of male WT DRG cultures, show a calcium response peak at 8 min upon application of MO (100μM), followed by a wash with Tyrode’s solution and another calcium response peak at 14 min upon application of KCl. C) Representative traces of male WT DRG cultures, show a calcium response peak at 8 min upon application of U50+MO, followed by a wash with Tyrode’s solution and another calcium response peak at 14 min upon application of KCl. D) One-Way ANOVA analysis revealed a significant treatment effect *F* _(2, 148)_ = 138.3 *P*<0.001 followed by Tukey’s *post-hoc* test. U50+MO significantly potentiates calcium release compared to MO alone (U50+MO vs. MO alone \*\**P* < 0.01). U50 treatment alone had no effect on the calcium response compared to MO alone and MO+U50 treatments (MO vs. U50 \*\*\**P*< 0.001, MO+U50 vs. U50 \*\*\**P*< 0.001). Data are presented as mean±SEM; n= 26-82/group. E) Representative traces of male Trpa1^−/−^ DRG cultures, show no calcium response peak at 4 min upon application of MO (100μM), followed by a wash with Tyrode’s solution and a calcium response peak at 9 min upon application of KCl. F) Representative traces of male Trpa1^−/−^ DRG cultures, show no calcium response peak at 4 min upon application of U50+MO, followed by a wash with Tyrode’s solution and a calcium response peak at 9 min upon application of KCl. G) MO vs. MO+U50 has no effect in Trpa1^−/−^ DRG cultures. Data are expressed as mean±SEM; n = 15-30/group.

## Discussion

Here we report that peripheral KOR activation increases noxious cold sensitivity in mice. We show that transcripts for KOR are present in the same cells as TRPA1 and TRPM8 in DRG. Furthermore, activation of KOR and TRPA1 together in cultured DRG potentiates calcium mobilization compared to activation of TRPA1 alone. This suggests that KOR are likely able to modulate noxious cold hypersensitivity through modification of TRPA1 in DRG; however, the intracellular mechanism between these two receptors has yet to be established.

KOR have long been considered promising targets for pain and itch relief due to their non-addictive profile (Porreca and Burks, 1983; Porreca et al., 1984; Shippenberg et al., 1988). However, the most significant limitation to targeting the KOR system has been the resultant negative affect, primarily mediated by the central activation of KOR (Horan and Porreca, 1993; Porreca et al., 1987), particularly in the nucleus accumbens (Al-Hasani et al., 2015; Massaly et al., 2019; Shippenberg et al., 1988). Recent evidence suggests that centrally- and peripherally-expressed KOR modulate different behaviors. For example, our group and others have shown that central KOR activation and upregulation modulates negative affect associated with peripheral nerve injury and inflammatory pain models (S. S. Liu et al., 2019; Massaly et al., 2019). Conversely, recent studies suggest peripherally-restricted KOR agonists selectively inhibit chemical pain and mechanical hypersensitivity associated with capsaicin-induced neurogenic inflammatory pain model and a surgical incision model, respectively (Snyder et al., 2018). Together these findings highlight the complex role of the KOR system in different pain and sensation modalities. As a result, this has prompted the investigation into the therapeutic potential of peripherally-expressed KOR (Beck et al., 2019; Snyder et al., 2018; Togashi et al., 2002). Progress has been constrained by the lack of peripherally restricted KOR compounds, as well as short-acting antagonists. However, more recently, there has been a focused effort to develop both short-acting reversible antagonists (Page et al., 2019) and peripherally restricted agonists (Barber et al., 1994; Beck et al., 2019; Olesen et al., 2013; Paton et al., 2020; Shaw et al., 1989; Suzuki et al., 2017) to understand the role of KOR in pain- and itch-related behaviors.

Thus far, in the itch field, Asimadoline (peripherally-restricted KOR agonist) has demonstrated efficacy in animal models of pruritus (Barber et al., 1994). The drug candidate is now in Phase 2 Proof-of-Concept clinical study to treat pruritus associated with atopic dermatitis by Tioga Pharmaceuticals (ClinicalTrials.gov, NCT01513161). TRK-820 (nalfurafine) manufactured by Toray Industries, Inc. is approved and used in Japan to treat uremic pruritus (Kumagai et al., 2010). JT09, a peripherally-restricted KOR agonist, has been shown to be as effective as morphine in alleviating pain without any sedative effects (Beck et al., 2019). CR845 is currently in phase II/III development to treat acute post-operative uremic pruritus and acute postoperative pain (ClinicalTrials.gov, NCT03998163) (Keppel Hesselink, 2017). The peptide is well-tolerated and is proven to be as effective as oxycodone in a human model of acute visceral pain (Arendt-Nielsen et al., 2009), without reports of dysphoria or hallucinations (Keppel Hesselink, 2017). In our current studies, we use an analogue of this compound, ff(nle)r-NH2 (Alleyne et al, in revision), to show that activation of peripheral KOR increases cold sensitivity in male and female mice, identifying an important role for the peripheral KOR system in noxious cold hypersensitivity.

Though little is known about the KOR system in cold sensation, it has been widely shown that the role of KOR in pain is dependent on the type of pain and sex. For example, in chronic pain states, KOR has been shown to induce negative affect (Liu et al., 2019; Massaly et al., 2019) that is absent following acute pain (Bagdas et al., 2016; Leitl et al., 2014b, 2014a). In both preclinical animal models and human imaging studies, KOR modulation of pain has been shown to be sex- dependent. In preclinical studies, intraplantar administration of U50 (100 μg/20μl) in males potentiated anti-hyperalgesic activity, as compared to females, suggesting a sex-dependent effect in the lateral sensitization rat model (Auh and Ro, 2012; Custodio-Patsey et al., 2020). Similarly administration of U50 increased tail withdrawal latency in male mice compared to female mice in tail withdrawal assay at 49 °C (Taylor et al., 2015). In addition, negative affect observed in peripheral nerve injury (PNI), neuropathic pain is potentiated in male mice alone following activation of KOR by U50, (Liu et al., 2019). Furthermore, the KOR-potentiated negative affect was blocked by JDTic (KOR antagonist) in male PNI mice, but not female PNI mice (Liu et al., 2019). Similarly negative affect associated in chronic inflammatory pain model was mitigated by JDTic in male mice (Liu et al., 2019). These preclinical findings show that the differential effects of sex not only directly impact pain processing, but also the negative affective state associated with the chronic pain. Clinically, positron emission tomography studies show that KOR receptor binding is higher in men than in women, especially in the anterior cingulate cortex, a region associated with pain affect (Vijay et al., 2018).

To fully evaluate sex as a biological variable, we investigated KOR-modulation of noxious cold hypersensitivity in both males and females. In this context, we report no significant differences except for the higher co-expression of *oprk1* and *trpm8* transcripts in males compared to females in DRGs. This is interesting as we also show that KOR-mediated cold hypersensitivity is only seen in females at 10°C (i.e., latency to jump on the cold plate at 10°C). Studies have evaluated the role of TRPM8 channels in mediating cold behavior at non-noxious temperatures 8°C −15°C (McKemy, 2007; Yang et al., 2020). Together these findings show that lower expression of TRPM8+KOR in females may play compensatory role in potentiating cold sensitivity at 10°C. Furthermore, the differential expression of TRPM8+KOR colocalization in DRGs between males and females has yet to be investigated in the presence of a conditional or global TRPM8 KO, but the sensitivity to noxious cold temperature upon activation of KOR seems to be minimal between males and females.

We also considered KOR’s somewhat controversial role in thermoregulation (Rawls and Benamar, 2011). At higher doses than used in this study (20 mg/kg; 40 mg/kg U50) activation of KOR has been shown to cause hypothermia in mice (Nemmani et al., 2001), and this is markedly influenced by tolerance developed by receptors upon repeated administration of KOR agonists (Milanés et al., 1991; Rawls et al., 2008; Von Voigtlander and Lewis, 1982). In contradiction, another study revealed that the 30 mg/kg dose of U50 did not affect rectal body temperature in mice (Itoh et al., 1993). Here we show that neither KOR agonists (U50 or ff(nle)r-NH2) alter body temperature in male and female mice.

To understand how activation of KOR mediates noxious cold hypersensitivity, we investigated TRPA1 and TRPM8 channels, both well known for their role in cold sensitivity (MacDonald et al., 2021; McKemy et al., 2002b; Palkar et al., 2015; Patapoutian et al., 2009). Interestingly, there has been targeted research exploring MOR, but not KOR, and TRP channel interactions (Shapovalov et al., 2013; Williams et al., 2013). In mice, spinal TRPA1 is shown to facilitate morphine-induced antinociception on a hot plate (Wei et al., 2016), suggesting a functional interaction between MOR and TRP channels in modulating thermal sensitivities. Furthermore, administration of opioids has been shown to distort thermal sensation; for example, some patients experience waves of warmth upon opioid administration (Chu et al., 2006), and drug withdrawal is often characterized by cold chills in combination with hyperalgesia (Pud et al., 2006). Similar information about KOR and TRP channels interactions are absent despite the fact that KOR is known to be involved in neuropathic pain (S. Liu et al., 2016; Navratilova et al., 2019; Xu et al., 2004).

Co-localization and electrophysiological studies have confirmed the presence of KOR on C- and A-fibers on DRG neurons (Ji et al., 1995; Zhang et al., 1998) expressing TRPV1, and calcitonin gene-related peptide (Snyder et al., 2018). Studies have shown that TRPM3 (Mucopilin 3), known to detect temperature and pain, interacts with the βγ subunits of the G-protein in the DRG to regulate calcium currents, implying the role of GPCR’s in calcium signaling (Badheka et al., 2015; Quallo et al., 2017). Similarly, MOR activation can inhibit the activity of TRPV1 via Gi/o proteins and the cAMP pathway indirectly effecting the capsaicin evoked calcium current in HEK293 cells, which shows how opioid receptors can mediate calcium activity via TRP channels (Endres-Becker et al., 2007; Vetter et al., 2006).

Interestingly, calcium imaging studies in human DRG neurons show a decrease in Ca^2+^ influx in the presence of the endogenous KOR ligand, dynorphin (Snyder et al., 2018). However, the calcium activity of KOR has not been evaluated in conjunction with TRP channels. We show that activation of KOR and TRPA1 together, in DRG, potentiates calcium release, as compared to activation of TRPA1 alone. This suggests that noxious cold hypersensitivity may be driven by peripheral activation of KOR that subsequently enhances the function of TRPA1, perhaps through receptor operation. Bautista et al. have shown similar mechanisms where Ca^2+^ released in response to inflammatory mediators act as a co-factor in activating TRPA1 (Bautista et al., 2006b, 2007).

Here our work elucidates the role of KOR action via TRPA1 channels potentiating calcium signaling which might underlie the cold hypersensitivity, conversely Snyder et al. elucidated the role of peripheral KOR activation in mitigation of chemical pain (Snyder et al., 2018). These findings together with recent data showing that cold sensing neurons mediate mechanical stimuli (MacDonald et al., 2021) suggest that these cold sensing neurons likely play a dual role in modulating noxious cold hypersensitivity and chemical pain. Exploration of the downstream signaling pathway mediating the calcium homeostasis via the KOR is warranted in order to understand the canonical/non-canonical pathway meditating KOR-induced noxious cold hypersensitivity.

Cold hypersensitivity is a chronic, debilitating, and poorly treated condition prevalent in neuropathic pain conditions such as multiple sclerosis, fibromyalgia, complex regional pain syndrome, and neuropathy following chemotherapy treatment. Our results identify a potential role for the KOR system in the mediation of noxious cold hypersensitivity. Here we show KOR’s role to be restricted to activation of peripheral KOR, which is encouraging and allows the study of this system without the aversive centrally-mediated side effects.

## Acknowledgments

We thank all the members of the Al-Hasani and McCall laboratories for helpful insight and discussion, especially to Gray B. Gereau, Marie C. Walicki, Gavin P. Schmitz, Kia M. Barclay, Lamley A. Lawson and Tori G. Collins for proficiently managing our lab’s mouse colony. We are thankful for technical assistance from the Gereau lab (WUSTL), Dr. Hongzhen Hu, (WUSTL), Dr. Steve Davidson (University of Cincinnati) and Dr. D.P. Mohapatra. Special thanks to Dr. Michael R. Bruchas and Dr. Gina M. Story for early support and pursuit of this line of work. We would also like to thank our administrative manager at the Center for Clinical Pharmacology, Jodi Maslin, for her support.

## Funding

NIH-R00DA038725 National Institute on Drug Abuse (R.A), NIH-R01NS117899 (J.G.M.), and the Rita Allen Foundation with help from the Open Philanthropy Projection (J.G.M.)

## DECLARATION OF INTERESTS

The authors declare no competing interests.

### Author contributions

Conceptualization, M.K.M, T.D.S., A.M.F., and R.A.; Methodology, M.K.M., L.V.T, J.G.M., and R.A.; Investigation, M.K.M., L.V.T, P.C., S.P., J.S.A., R.A.H., and R.A. Manuscript preparation, M.K.M., J.G.M., and R.A.; Funding acquisition, R.A.; Supervision, J.P.M., J.G.M., and R.A.; Project administration, R.A.

## Supplementary figures

**Figure S1.**
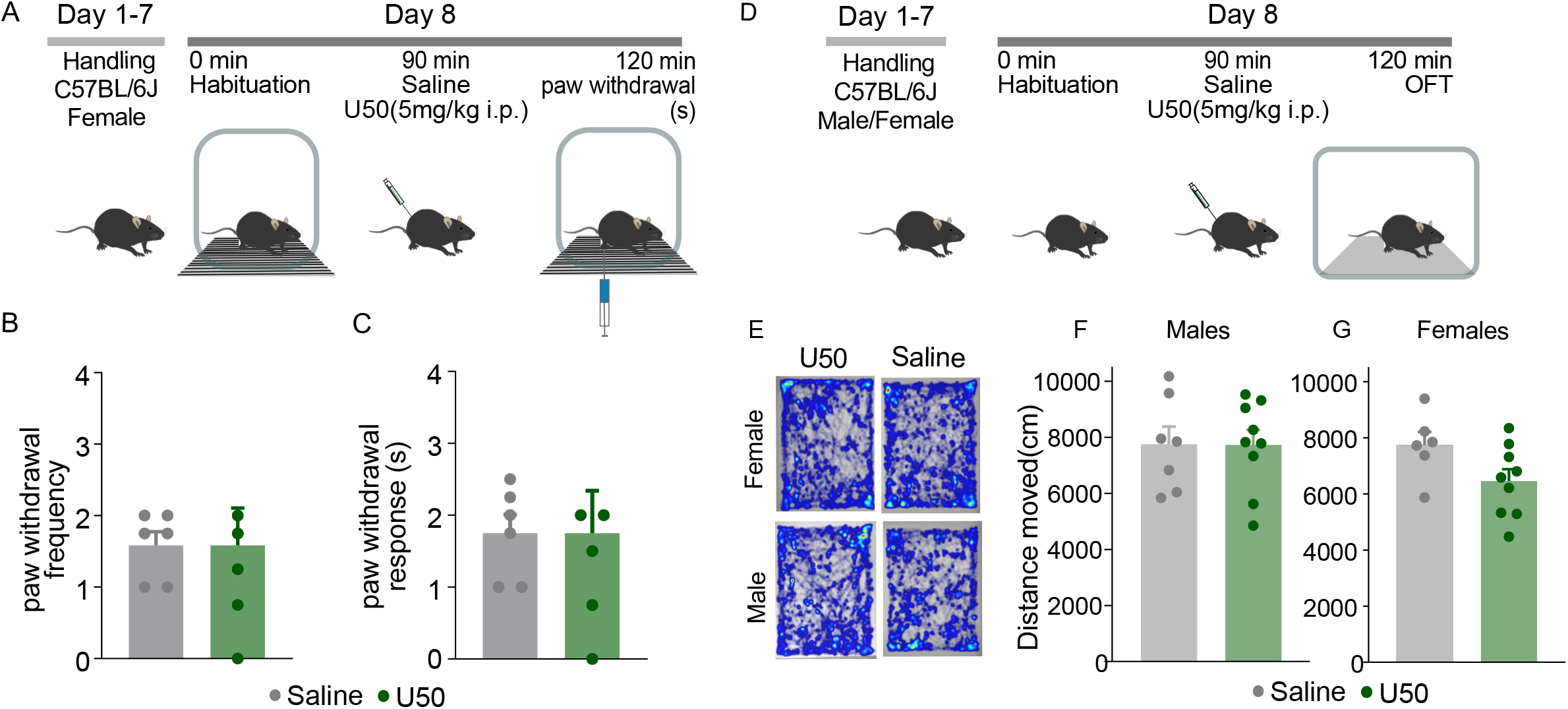
A) Outline of the acetone evaporation test experimental procedure. Timeline of the systemic injections of U50, followed by habituation and acetone evaporation assay. B) U50 had no effect on acetone-evoked aversive behavior, as compared to saline, in females. The acetone-evoked aversive behavior was measured by quantifying the nocifensive responses (scoring the paw withdrawal frequency). Data are expressed as mean± SEM; n=5-6/group. C) U50 had no effect on acetone-evoked aversive behavior, as compared to saline, in females. The acetone-evoked aversive behaviour was measured by the duration of nocifensive response (scoring the latency of paw withdrawal response). Data are expressed as mean± SEM; n=5-6/group. D) Outline of the experimental procedure. Timeline of the open field test (OFT) assay, habituation followed systemic injections of saline or U50, and performing OFT. E) Representative heat maps of saline and U50 treatment groups in OFT. F) U50 had no sedative effect in the open field test in males. Data are expressed as mean± SEM; n=6-9/group. G) U50 had no sedative effect in the open field test in females. Data are expressed as mean± SEM; n=6-9/group.

**Figure S2.**
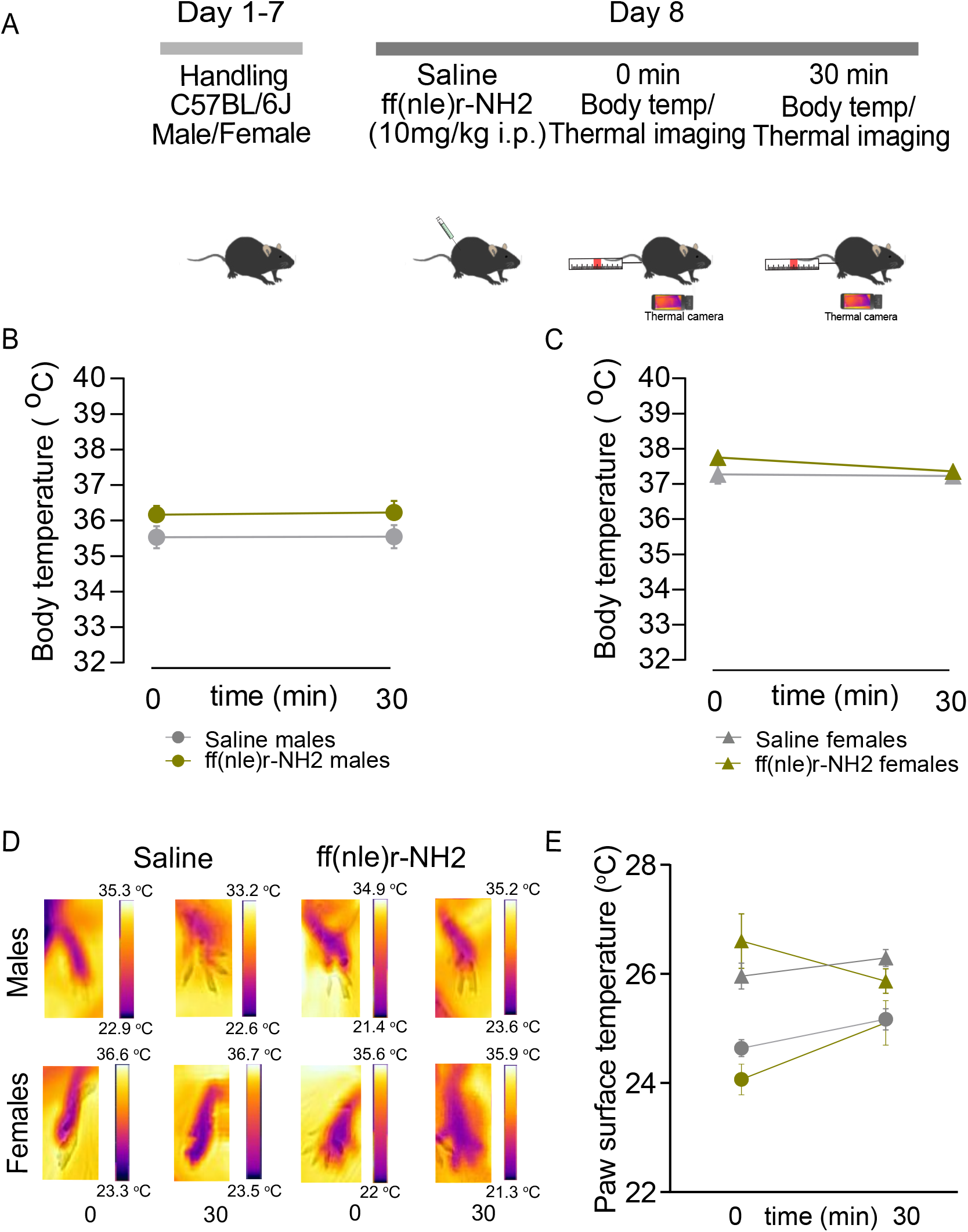
A) Outline of the experimental procedure. Timeline of the assay, wherein mice are habituated, baseline rectal temperature is measured along with thermal imaging of paw surface temperatures, followed by systemic injections of saline or ff(nle)r-NH2 and post-treatment temperatures were recorded. B) Peripheral KOR agonist ff(nle)r-NH2 did not alter core body temperature 30 min post drug administration in males. Data are expressed as mean±SEM; n=6-10/group. C) Peripheral KOR agonist ff(nle)r-NH2 did not affect core body temperature 30 mins post drug administration in females. Data are expressed as mean±SEM; n=6-10/group. D) Representative infrared images from right hind paws of a male and female mouse using FLIR infrared camera pre-and post-saline or ff(nle)r-NH2 adminstration. E) KOR agonist ff(nle)r-NH2 did not alter paw surface temperatures in males and females. Data are expressed as mean±SEM; n=4/group.

## Notes

### Competing Interest Statement

The authors have declared no competing interest.

### Summary of Updates

figures 3 and 4 have been reformatted and the text has been revised. Figure 3 has more detailed co-localization analysis, Figure 4 added a U50 only condition in the TRPA1 knockout group. Added second supplementary figure showing body temperature and thermal imaging following administration of peripheral compound.

